# Structural Capacitance in Protein Evolution and Human Diseases

**DOI:** 10.1101/269613

**Authors:** Chen Li, Liah V. T. Clark, Rory Zhang, Benjamin T. Porebski, Julia M. McCoey, Natalie A. Borg, Geoffrey I. Webb, Itamar Kass, Malcolm Buckle, Jiangning Song, Adrian Woolfson, Ashley M. Buckle

## Abstract

Canonical mechanisms of protein evolution include the duplication and diversification of pre-existing folds through genetic alterations that include point mutations, insertions, deletions, and copy number amplifications, as well as post-translational modifications that modify processes such as folding efficiency and cellular localization. Following a survey of the human mutation database, we have identified an additional mechanism, that we term ‘structural capacitance’, which results in the *de novo* generation of microstructure in previously disordered regions. We suggest that the potential for structural capacitance confers select proteins with the capacity to evolve over rapid timescales, facilitating saltatory evolution as opoposed to exclusively canonical Darwinian mechanisms. Our results implicate the elements of protein microstructure generated by this distinct mechanism in the pathogenesis of a wide variety of human diseases. The benefits of rapidly furnishing the potential for evolutionary change conferred by structural capacitance are consequently counterbalanced by this accompanying risk, with the extent of this determined by the host immune system. The phenomenon of structural capacitance has implications ranging from the ancestral diversification of protein folds to the engineering of synthetic proteins with enhanced evolvability.

## Introduction

Canonical protein evolution is achieved through the utilization of an array of different genetic mechanisms, including point mutations, recombination, translocations, and duplication. Alterations to the protein-encoding elements of genes are accompanied by modifications to non-coding control elements, epigenetic changes and post-translational mechanisms that refine the kinetics, spatial localization, synthesis, folding efficiency, and other aspects of protein synthesis and dynamics. The functional and morphological diversity at the protein, cellular, and organismal level is achieved through the utilization of a combination of such mechanisms. This repertoire of available mechanisms for the implemention of evolutionary change enables adaptations to be furnished at the appropriate level, and allows proteins to efficiently navigate their function spaces to identify optimal phenotypic solutions.

Whereas classic genetic modifications are incremental, more complex mechanisms, such as the phenomenon of genetic capacitance mediated by heat shock proteins like Hsp90, buffer the impact of polymorphisms that may in isolation be maladaptive, allowing their phenotype to remain hidden and offering the possibility for ‘saltatory’ evolutionary change [1]. This enables expansive regions of structural and functional space to be navigated in a single step, releasing phenotypes from local optima and allowing new functions and morphologies to evolve over rapid timescales along routes requiring the simultaneous presence of multiple genetic alterations. The existence of such mechanisms enables evolutionary landscapes [1] to be navigated more efficiently, as searches may be extended beyond the local optima of rugged peaks, so as to reach out across otherwise un-navigable regions of sequence space to identify distant, and identify potentially more adaptive optima. The capacity for this type of accelerated evolution is especially important in environments chracterized by high uncertainty, where a capacity for furnishing rapid and efficient evolutionary change is a prerequisite for survival. Interestingly, there is emerging evidence for profound protein structural changes induced by so-called “hopeful monster” mutations [2–4].

Many proteins, and most notably those involved in cellular regulation and signaling [5, 6] contain significant regions of disorder, and belong to the class of Intrinsically Disordered Proteins (IDPs) [7]. IDPs can undergo disorder-order transitions, typically upon binding another protein or DNA (coupled folding and binding [8–11]). The energy landscapes of IDPs are typically rugged, featuring a continuum of conformational states that enables interaction with other molecules via both conformational selection and induced fit [8, 12, 13].

In order to establish whether point mutations within the regions encoding disordered regions may result in microstructuralization and generate nascent microstructural elements that may form substrates for evolution or result in adaptive alterations to protein function, we performed a survey of the human mutation database [14]. Specifically, we performed a bioinformatic analysis to identify mutations predicted to generate localized regions of microstructure in previously disordered regions of target proteins. We report here a new mechanism of protein evolution, which we term ‘structural capacitance’, whereby structural and functional changes at the level of individual proteins may be achieved through the introduction of point mutations influencing key nucleating amino acids located in regions of structural disorder. Once mutated, these residues are predicted to generate new microstructural elements in previously disordered regions that are functionally distinct from the parent fold. These findings have broad implications both for the accelerated non-canonical evolution of protein folds, and for the pathogenesis of human diseases.

## Results

### Order-disorder transitions associated with mutations in human proteins

We interrogated the human polymorphisms and disease mutations dataset [14], and compiled a dataset of 68,383 unique human mutations (excluding the ‘unclassified’ mutations; see ‘Materials and Methods’ for more details), comprising 28,662 human disease mutations, and 39,721 polymorphisms. We then applied standard algorithms for disorder prediction to every mutation and polymorphism in the dataset. A predictor voting strategy was employed to determine the prediction outcome for each mutation. As there are multiple predictors for protein disorder, the residues were deemed to be located within disordered regions if the number of predictors assigning residues to disordered regions were equal to or larger than the number of predictors assigning the residues to ordered regions. Four types of structural transitions were defined: Disorder-to-Order (D**→**O), Order-to-Disorder (O**→**D), Disorder-to-Disorder (D**→**D) and Order-to-Order (O**→**O) (Table S1).

We next interrogated the subset of proteins containing D**→**O predicted mutations imposing the following selection criteria: (1) mutations located within disordered regions ≥30 amino acids in length (termed “Long Disordered Regions” (LDRs), consistent with other studies [15–19]); (2) LDRs not predicted to be in transmembrane domains and (3) proteins with LDRs lacking experimentally determined identical or homologous structures. From a BLAST search against the protein databank (PDB), 231 out of 1,337 (17.3%) proteins do not currently have experimentally determined structures or homologues. Among 1,731 proteins containing predicted D**→**O mutations, we identified 152 point mutations within LDRs from a total of 133 proteins (Table S2). The workflow is detailed in Figure S1.

In order to determine whether any of the mutation sites are located in functionally-relevant regions, we cross-referenced disease and non-disease mutations in D**→**O, O**→**D, D**→**D and O**→**O transitions with the Eukaryotic Linear Motif (ELM) database [20]. Eukaryotic linear motifs (ELMs) are predominantly functional modules found in intrinsically disordered regions in eukaryotic proteins [21]. All ELMs listed have been experimentally verified (i.e., annotated with experimental evidence showing that the ELM is involved in a functionally-relevant interaction). The number of mutations found in ELMs were: 13/1,731 (0.75%), 76/13,876 (0.55%), 75/51,317 (0.15%) and 3/1,459 (0.21%) for D**→**O, D**→**D, O**→**O and O**→**D mutations respectively (Tables S3-S6). The number of identified motifs for D**→**O mutations is much smaller than for D**→**D and O**→**O, however, this most likely reflects the relative scarcity of D**→**O mutations. For all four transitions, more disease mutations than non-disease mutations/polymorphisms were found in ELMs according to the one-tailed Fisher exact test (8 *vs* 5 for D**→**O disease-associated mutations *vs* polymorphisms (*p*-value = 0.02); 3 *vs* 0 for O**→**D disease-associated mutations *vs* polymorphisms (*p*-value = 0.09); 46 *vs* 30 for D**→**D disease-associated mutations *vs* polymorphisms (*p*-value < 0.00001); 45 *vs* 30 for O**→**O disease-associated mutations *vs* polymorphisms (*p*-value = 0.03). Overall, therefore, we predict that only a relatively small fraction of the identified D**→**O mutations are part of functionally-relevant interactions with other proteins. To further investigate the numbers of mutations that are located in the annotated Pfam domains, we mapped the mutations of four transitions to Pfam database [12]. The numbers of mutations found in Pfam domains were: 741/1,731 (42.8%; 308 *vs* 433 for disease-associated mutations *vs* polymorphisms), 3,538/13,876 (25.5%; 1,333 *vs* 2,205 for disease-associated mutations *vs* polymorphisms), 800/1,459 (54.8%; 474 *vs* 326 for disease-associated mutations *vs* polymorphisms) and 35,521/51,317 (69.2%; 19,500 *vs* 16,021 for disease-associated mutations *vs* polymorphisms) for D**→**O, D**→**D, O**→**D and O**→**O, respectively. The mapping results for D**→**O and O**→**D transitions are shown in Tables S7 and S8, respectively. On average, approximately half of the mutations were located in Pfam annotated domains.

### Identification of Structural Capacitance Elements (SCEs)

We propose that the identified LDRs (Table S2) represent a new class of genetic element, which we have termed ‘Structural Capacitance Element’ (SCE). The D**→**O mutations identified represent examples of order-inducing substitutions that introduce new microstructure into the parent fold that may confer new functions, or refine existing ones, but which may, in some cases, be of pathogenic significance. There are 21 mutations involving cysteine residues identified within 21 proteins (all involving a single mutation to cysteine; Table S2). 0.75% of the identified D**→**O mutations are predicted to be associated with ELMs (Tables S3), indicating that these substitutions are unlikely to interfere with known interactions, resulting instead in the potential for functionality through the generation of new microstructures. The mechanism appears to be initiated by point mutations that change hydrophilic nucleating residues to hydrophobic ones (Figure 1A). It remains to be seen whether the codons encoding these residues constitute ‘hotspot’ regions within the human genome, and whether there is codon bias at these positions. To investigate the possible effects of D**→**O mutations on proteolytic processing, we mapped these 152 D**→**O disease-associated mutations (Table S2) to a list of experimentally verified proteolytic cleavage sites from the MEROPS database [22]. We identified 67 cleavage sites located within 15 residues of the disease-causing D**→**O mutation sites belonging to 36 proteins (Table S9).

**Fig. 1.**
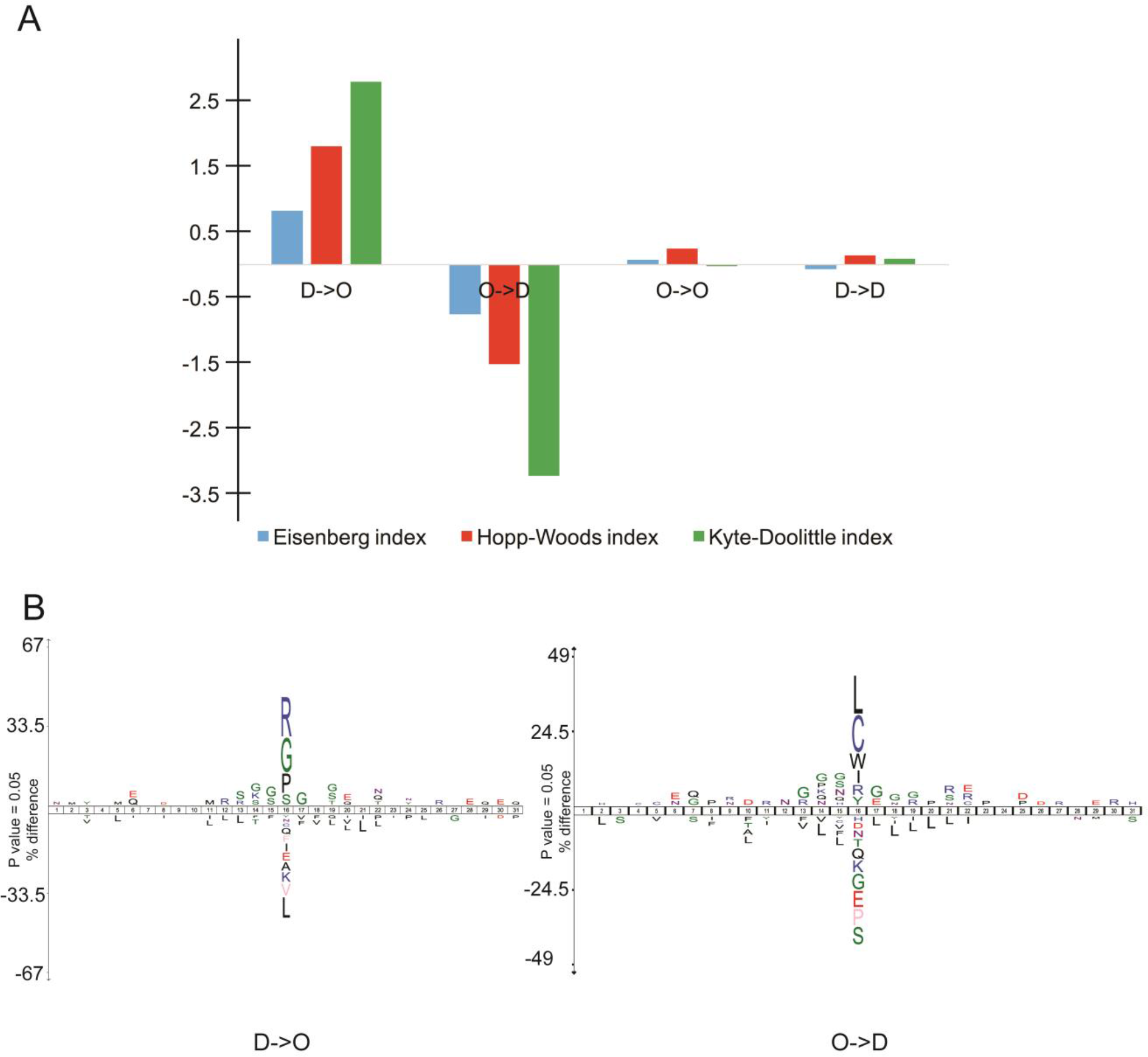
**(A)** Mean hydrophobicity changes for disease-causing mutations for four different classes of structure-altering mutations predicted using multiple sequence-based predictors for intrinsically disordered regions [23] (e.g., Disorder-to-Order is denoted as D**→**O, Order-to-Disorder as O**→**D). Bars are shown for the three different hydrophobicity indices used: Eisenberg hydrophobicity index [24] (Blue), Hopp-Woods hydrophilicity index [25] (Red) and Kyte-Doolittle hydropathy index [26] (Green); **(B)** IceLogo [27] charts showing the residue conservation around the disease-causing mutation site against a reference set (human Swiss-Prot proteome) for D**→**O and O**→**D transitions with wild-type residue in the central position. Amino acids residues on top of the *x* axis are significantly conserved, while those underneath it are non-preferred or unfavoured according to the reference set.

The types of mutations in each class (D**→**O, O**→**D, D**→**D and O**→**O) appear to be non-random (Table S10). For all documented D**→**O disease mutations, arginine is the most frequently mutated amino acid (Figure 1B). The most common classes of disease mutation for D**→**O and O**→**D transitions are R**→**W (59, 11.50%) and L**→**P (108, 16.62%), respectively (Tables S10A and S10B). For O**→**O and D**→**D transitions, the mutation patterns are more evenly distributed (Tables S10C and S10D). For non-disease mutations, the most common type is P**→**L (153, 12.56%) for D**→**O and L**→**P (63, 7.79%) for O**→**D (Tables S10A and S10B). This is consistent with a recent comparison of mutation frequencies in intrinsically disordered regions of proteins in both disease and non-disease datasets that highlights the previously unappreciated role of mutations in disordered regions [11]. An example of the predicted local increase in ordering due to D**→**O mutations in both disease and non-disease datasets is shown in Figure 2A and B.

**Fig. 2.**
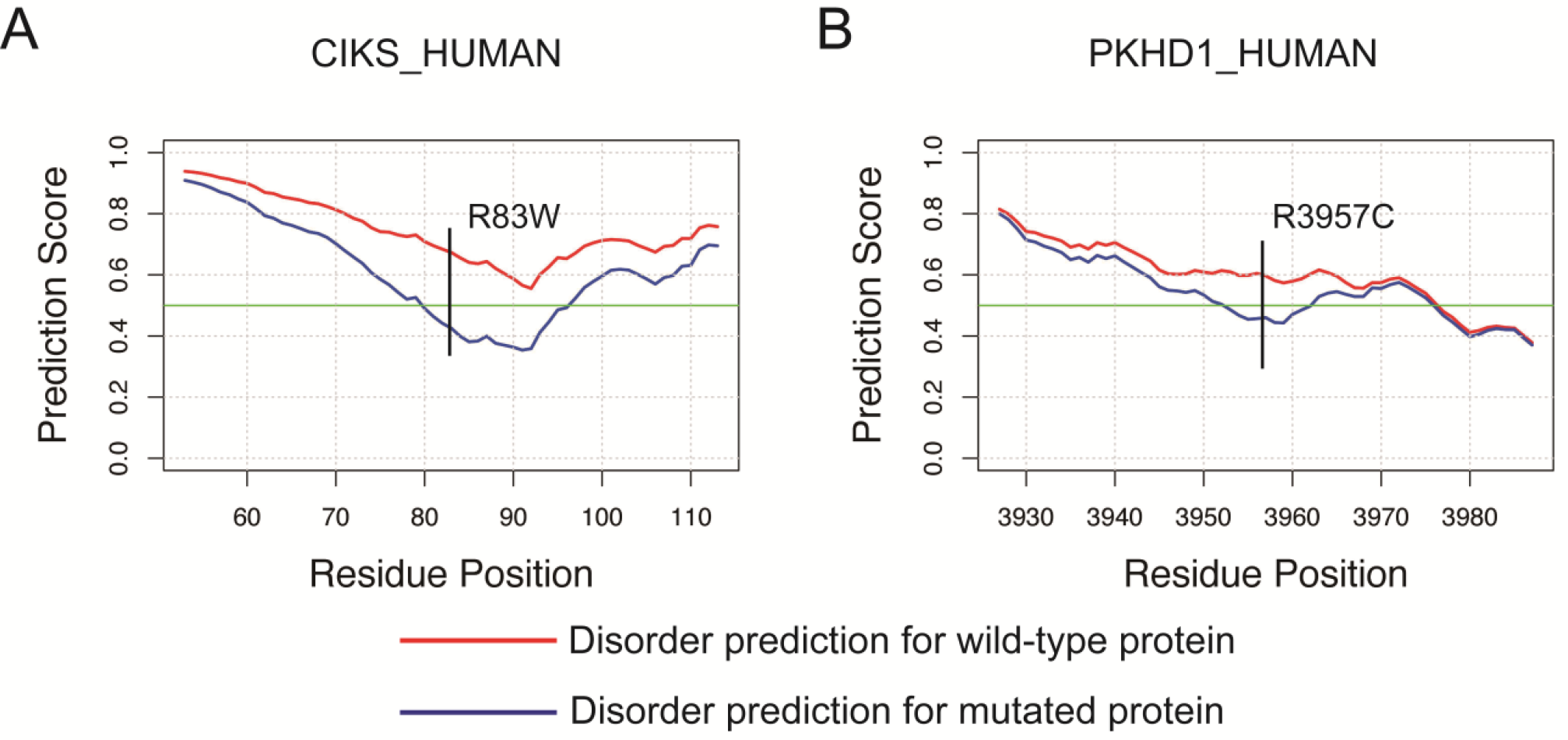
Predicted disorder score change according to VSL2B for **(A)** Adapter protein CIKS (UniProt ID: O43734 / CIKS_HUMAN; Polymorphism); **(B)** Fibrocystin (UniProt ID: P08F94 / PKHD1_HUMAN; Polycystic kidney disease; autosomal recessive (ARPKD); [MIM:263200]).

Given the variability in the predictions among the four predictors tested, which necessitated a majority voting approach, we looked for experimental evidence suggesting that the predicted regions were disordered. Accordingly, we cross-referenced our human disease mutations and polymorphisms dataset against DisProt [28], a database providing experimentally verified disordered regions of proteins. For the resulting matches, we applied four protein disordered region predictors to predict the structural changes following mutation events, using majority voting. From our original dataset of 76,608 mutations, 409 mutations (0.5%) are found in DisProt to map to experimentally validated disordered regions. Disorder prediction using majority voting predicts that 132 mutations in these regions result in a D**→**O structural transition (Table S11). The protein context information showing the distribution of protein domains, post-translational modification sites and mutations of protein entries in Table S11 are shown in Table S12. Examples showing increase in ordering according to DynaMine based on the DisProt dataset are shown in Figure S2.

## Discussion

The association between disease phenotypes and the *de novo* formation of protein microstructural elements represents a novel paradigm for understanding the origins of select human diseases, which has principally been focused on loss-of-structure and accompanying loss-of-function. One of the best studied examples is the tumor suppressor p53, which is inactivated following somatic mutation events in a range of human cancers [29]. Most p53 mutations are loss-of-function mutations impacting the DNA-binding domain through interference with p53-DNA contacts or structural destabilization [30]. It is, however, known that IDPs play a role in cellular regulation and signaling [5, 6]. Indeed structural changes in disordered proteins have been implicated in disease processes, with evidence for D**→**O transitions triggered by disease-causing mutations in functionally important regions (such as regions mediating protein-protein recognition via coupled folding and binding, and DNA-binding [11] including p53-DNA binding [10]).

We propose that in addition to canonical loss-of-function mutations, disease-causing mutations may result in gain-of-function through the binary activation of cryptic ‘structural capacitance elements’ (SCEs, Figure 3). Our findings are in broad agreement with a recent analysis of disease-causing mutations in disordered regions, which demonstrated that a significant number of D**→**O mutations are predicted to disrupt protein function[11]. However, here we distinguish mutations that disrupt disorder-based functional properties from those that induce microstructuralization and accompanying gain-of-function through the phenomenon of structural capacitance. It is possible that some of the D**→**O mutations we have identified may induce pathological changes through disrupting known associations with interaction partners, for example via premature microstructuralization. The lack of evidence for functional interactions and an analysis of ELMs [31] nevertheless suggest that few residues undergoing D**→**O mutations form part of an interaction with another protein (Table S3).

**Fig. 3.**
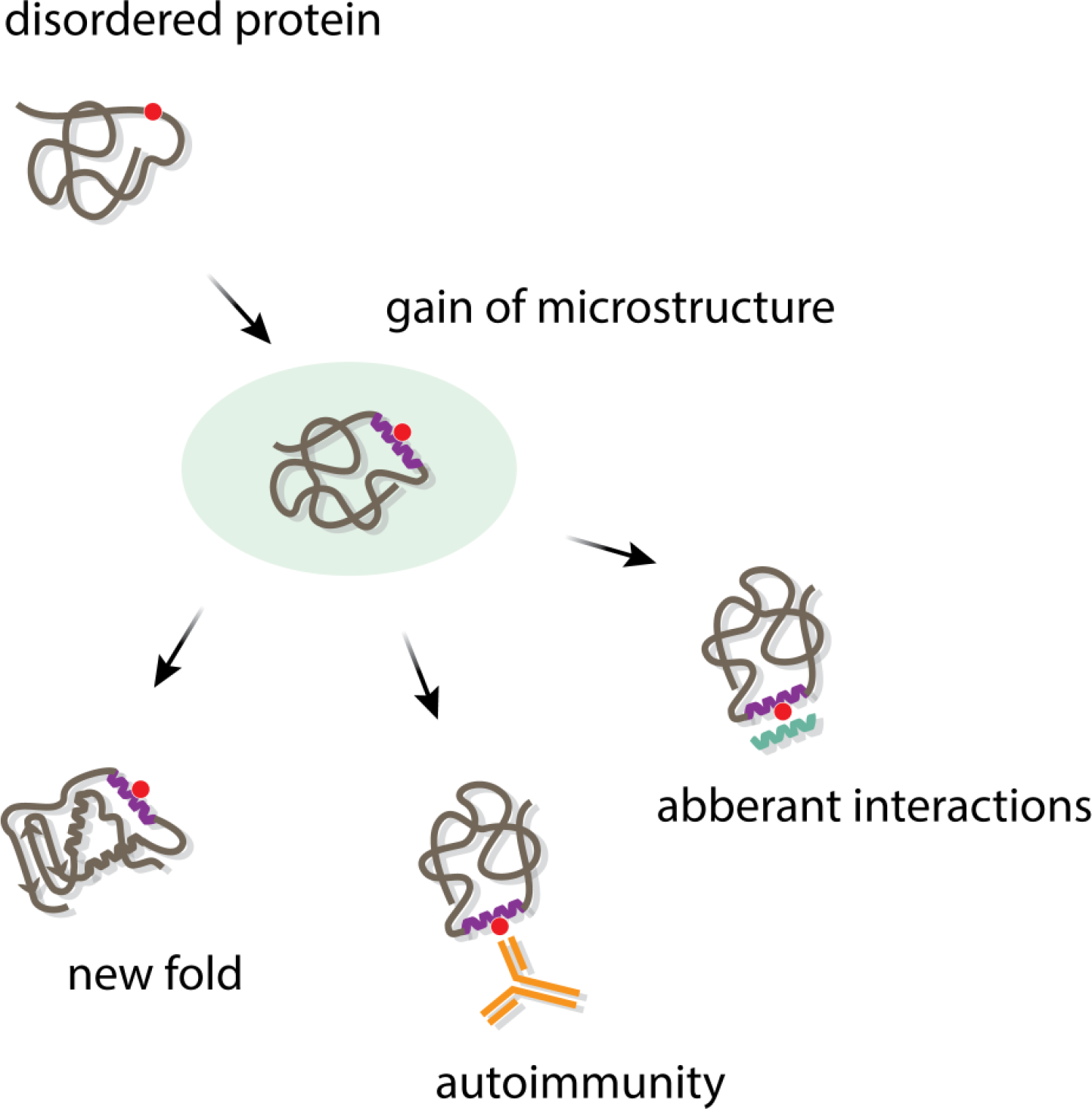
Disease-causing mutations may result in gain-of-function through the mechanism of structural capacitance. A D**→**O mutation (red circle) in a disordered protein results in the generation of local microstructure, or ‘structural capacitance element (SCE)’ (purple helix). This may be a key nucleating factor in the microevolution of a new adaptive fold, but may also generate inappropriate pathological interactions, through the triggering of inflammatory and autoimmune responses. Aberrant interactions may also promote other pathogenic processes such as aggregate formation, which may result in the formation of pathogenic fibrils.

In contrast to loss-of-function through loss-of-structure in canonical disease-causing mutations[29, 30], the complementary phenomenon of gain-of-structure and function through the introduction of microstructuralization into disordered or unstructured regions of proteins remains undescribed. The characterization of these changes is challenging due to the technical hurdles associated with resolving the structural properties of a structurally heterogeneous disordered population. Nevertheless, experimental evidence to support the pathogenic relevance of D**→**O transitions resulting from disease-associated mutations has been described in several proteins [32–34].

The phenomenon of structural capacitance has significant implications for protein evolution and for the diversification of organismal form and function over evolutionary time, and is likely to augment other known mechanisms of evolutionary modification such as genetic capacitance [1] and “cryptic’ genetic variation [35] that offer the potential for explorations of protein fold and morphological space over compressed timescales. Proteins with high flexibility and dynamics may have a greater intrinsic structural capacitance, increasing their evolvability, and allowing for the more rapid evolution of new folds [36]. The evolution of primordial proteins may have involved either the co-evolution of folds and functions through conformational selection from a repertoire of disordered polypeptides, or the emergence of secondary structure elements followed by the evolution of fully folded proteins [36]. Both scenarios, however, require the prior formation of local structure from an essentially random and disordered population. This provides a conundrum, as how would classic Darwinian evolution proceed in the absence of pre-existing seed structures and functions [37]? We suggest that the microstructuralization necessary for founder events in protein folds may be furnished by structural capacitance (Figure 3). In this mechanism, SCEs are localized regions of disorder retained within protein structures, and controlled by key capacitance residues that confer the potential to generate new microstructural elements that modify the evolvability of the fold. Structural capacitance generates the potential for micro-structural change, which helps buffer organisms against the vagaries of an uncertain future by furnishing adaptive solutions.

In the co-evolutionary model of protein fold and function, microstructuralization of conformationally diverse protein species within a population complements the conformational selection by ligands through the stabilization of functional conformers. The notion that functional selection may occur through the binding of small molecules is compelling, but this process may not inevitably require the prior formation of significant structural scaffolds and could proceed from a relatively small nucleus of structure. Recent work suggests that ligand-binding features arise from the physical and geometric properties of proteins, with structures serving as a feedstock for evolution [38]. This is consistent with findings from directed evolution studies that demonstrate the acquisition of function following only a few prior rounds of selection [39, 40]. Structural capacitance may provide the key nucleation event for the formation of a feedstock of molecules with weak functional activity that have the capacity to be fine-tuned and the potential to generate the specificity, high affinity binding and selectivity characteristic of modern enzymes.

In the alternative scenario where the evolution of the fold mimics the folding pathway and the ancestral progenitors of modern folds resemble folding intermediates [36], structural capacitance may introduce reversibility into evolutionary processes in a manner distinct from the ratchet-like and often irreversible mechanism of canonical incremental evolution. The energy landscape of an unstructured or disordered protein may be considered relatively flat with ruggedness depending on hydrophobicity and stereochemistry (Figure 4A). D**→**O mutations with SCEs that induce microstructuralization may induce small impressions, or high-altitude “fissures” into such geographical landscapes (Figure 4B). Microstructuralization is reversible, allowing rapid conformational transitions and landscape exploration, and minimizes sequestration in dead-end, local minima. The most powerful structural capacitance is predicted to be located in landscapes where D**→**O mutations, acting as reversible “binary switches”, are expected to have the most significant impact through introducing bias from one lake to another, or transitioning it into a stable ‘valley’. Canonical gradualistic evolution may then optimize the funnel characteristics of the energy landscape (Figure 4C). These features of structural capacitance would allow for rapid fold generation and may have facilitated some of the major transitions in organismal evolution, complementing and potentiating gradualistic modifications to pre-existing folds. Although there are notable exceptions (serpins, for example [41, 42]), proteins with highly evolved functions are generally situated in deep energy wells at the bottom of the folding landscape, or funnel, preventing major structural changes. Highly optimized active site architectures represent an irreversible evolutionary ‘ratchet’ that may limit the evolutionary adaptability of a fold because non-functional mutants are strongly selected against. Structural capacitance may circumvent such limitations, thereby providing a mechanism for evolvability, and could be exploited for the engineering of artificial proteins with an enhanced capacity for plasticity through micro-evolutionary change [43, 44]. Furthermore, such a mechanism might offer some molecular insights into the relationship between cryptic genetic variation [35] and protein evolvability – microstructuralisation within SCEs may enable “pre-adapted” phenotypes that confer selective advantages when new selection pressures emerge [45].

**Fig. 4.**
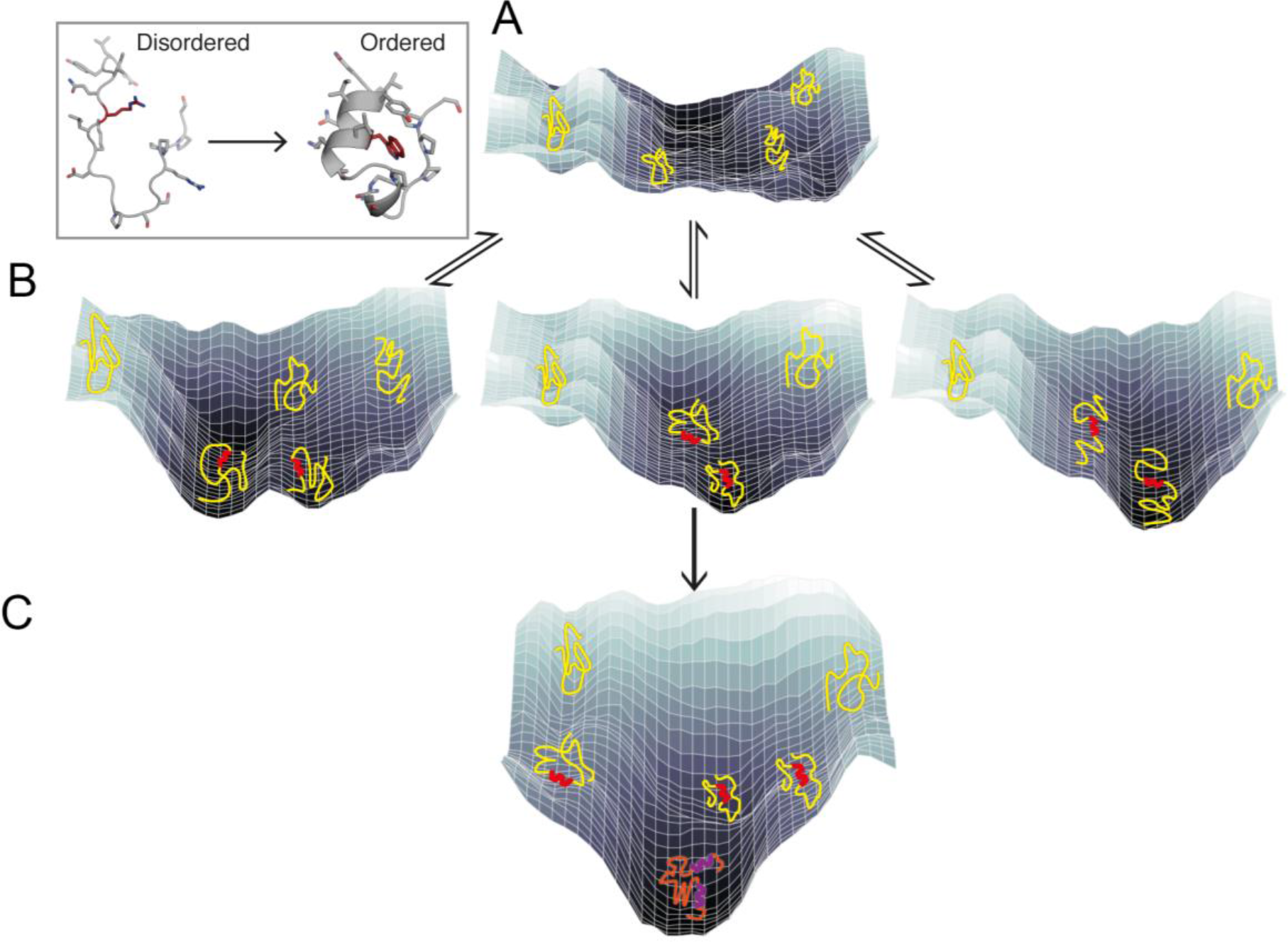
Structural capacitance and folding energy landscapes. **(A)** Flat, featureless energy landscape of a disordered protein; **(B)** D**→**O mutations in SCEs induce microstructure and small impressions, allowing conformational transitions and landscape exploration; **(C)** Canonical incremental evolution optimizes the folding funnel to create new fold. The inset depicts an R**→**W mutation (sticks) inducing helix formation and local structuralization, based upon the Trp-cage protein TC5b [46]. Hydrophobic clusters centered around tryptophan are common in several small natural folds [47, 48].

Protein folding may nucleate via relatively few key residues, which are typically hydrophobic [49]. Although the energy landscape in which IDPs bind to targets is likely to be relatively complex, experimental evidence supports conformational selection of secondary structure formation, followed by induced fit following binding and completion of global folding [8, 50, 51]. Our finding that O**→**D and D**→**O transitions are predominantly associated with mutations to and from proline, respectively (Tables S10A and S10B), is consistent with the demonstrated helix-disrupting properties of proline [52, 53]. It is intriguing that 59 (11.5%) of the disease-causing and 80 (6.6%) of non-disease D**→**O mutations reported involve a mutation to tryptophan. Hydrophobic clusters centered around tryptophan are common in several small natural folds [47, 48] most notably the Trp-cage fold, which is one of the smallest model proteins of just 20 amino acid residues and folds spontaneously into a stable 3D structure within ~4 μs, featuring a hydrophobic core formed around a central tryptophan residue [54, 55]. Along with evidence of residual structure due to hydrophobic collapse around the central tryptophan in the unfolded state of Trp-cage protein TC5b [46], our findings are consistent with tryptophan residues playing an important role in the generation of microstructure from disorder (Figure 4 inset).

We propose that the phenomenon of structural capacitance may be extended to a more general framework encompassing the roles of unstructured proteins (Figure 5). Approximately 40% of the human proteome is intrinsically disordered [56], but the selection advantage this provides has not been clearly defined. This reservoir of unstructured and highly dynamic protein sequence is highly evolvable for two reasons. First, via the now accepted mechanism of coupled folding and binding (reviewed in [9]) it may act in concert to engage a broad range of binding partners. This is consistent with the compelling evidence for the role of IDPs in signalling and interaction network hubs [27, 29, 56]. Second, as described in this work, D**→**O mutations in SCEs may generate highly evolvable species of conformations with microstructure over rapid timescales that facilitate the evolution of new folds. Both involve a D**→**O transition, whereby the information for folding is stored in the unstructured protein ensemble. This ensemble is highly evolvable and acts as a ‘structural capacitor’. Release of folding information through microstructuralization is achieved by the binding of a structured physiological partner, as is the case for coupled folding-and-binding of IDPs, or as the result of a D**→**O mutation. This concept encapsulates and extends the ‘dormant foldon’ hypothesis [57]. Furthermore, structural capacitance is compatible with the concept of early peptide-world ‘foldamers’ [58] as well as Dayhoff’s, and more recent hypotheses of early protein evolution driven by oligomerisation-duplication-fusion events of short peptides [59, 60].

**Fig. 5.**
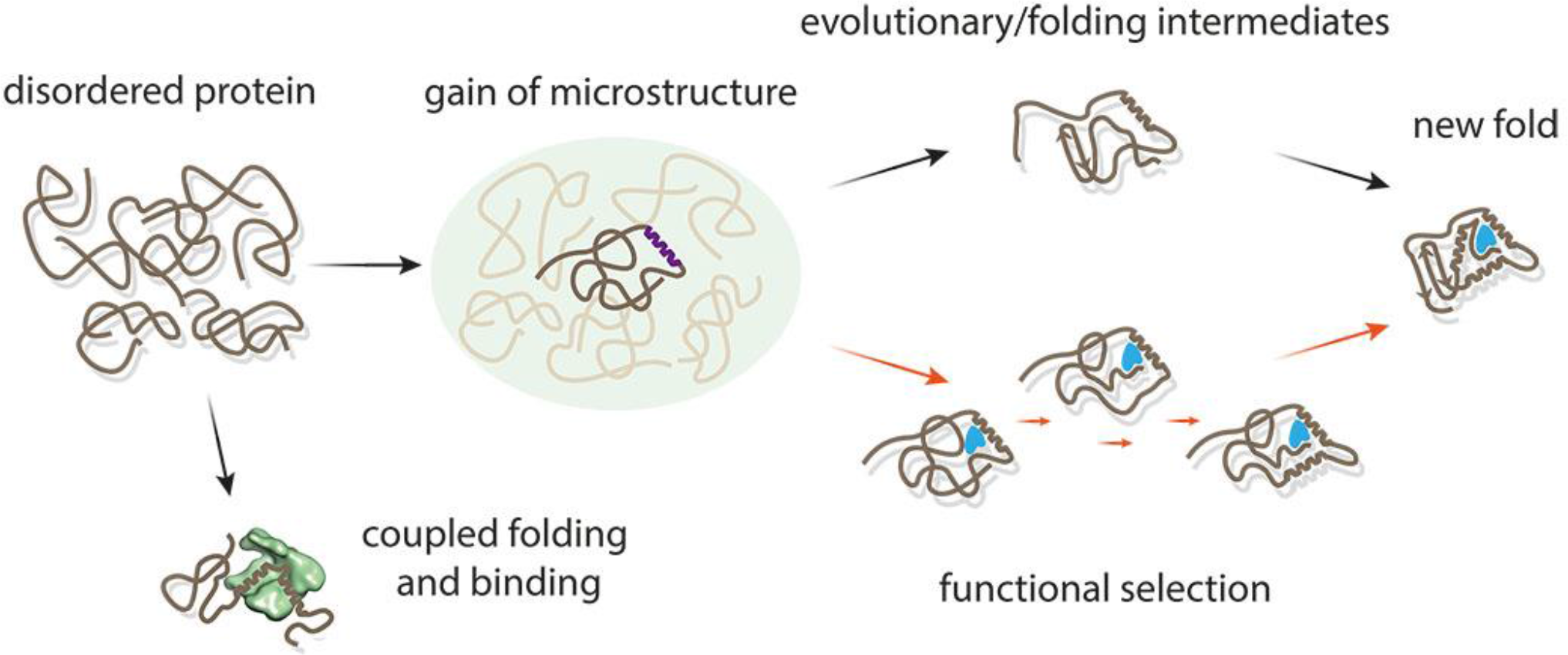
Structural capacitance incorporating the concepts of classic incremental Darwinian evolution, dynamism, evolvability, and structural diversity, provides a potential basis for the generation of microstructure from randomness. The ‘capacitor’ comprises a random ensemble of disordered protein conformations harboring SCEs. These may undergo ordering through either coupled folding-upon-binding of a partner (bottom left) or the fixation of a D**→**O mutation, that ‘discharges’ the folding information stored within its sequence. A D**→**O mutation may bias the random population towards intermittent microstructure, creating a highly evolvable feedstock for subsequent step-wise evolution through random mutation. This process may proceed by selection through either function or folding to produce a new fold.

### Further implications and hypotheses

#### Cellular proteostasis

Microstructuralization through D**→**O mutations has the potential to alter cellular proteostasis. Conventionally, the effect of mutations on protein degradation and proteostasis has been investigated in the context of O**→**D mutations; in the scenario of D**→**O mutations the opposite may be observed, with the acquisition of microstructure or ordering within a protein having the potential to slow the kinetics of its degradation and increase its longevity, perhaps eventually leading to the accumulation of toxic species. Previously exposed lysines that may be the target of a ubiquitin degradation signal may become concealed by the acquisition of microstructure and subsequent promiscuous formation of aberrant interactions, or perhaps self-assembly, decreasing proteasome-mediated degradation [61]. This scenario may ultimately promote the accumulation of toxic aggregates that overload the proteasomal processing capacity, with well-established implications for a wide range of pathologies (reviewed in [62]). Our analysis of the effects of D**→**O mutations on proteolytic processing is consistent with this hypothesis (Table S9).

#### Viral adaptation

The high mutational rate of viruses is a well-characterized phenomenon that allows for adaptation to rapidly changing environments. Recent reports implicate structural disorder in viral adaptation[18], and demonstrate how mutations in disordered regions may promote neostructuralization and accompanying phenotypic divergence[63, 64]. Structural capacitance within disordered ensembles of viral proteins may represent a powerful mechanism for enhanced pathogenicity through facilitating the rapid acquisition of microstructure and an accompanying improvement in the ability to interact with host proteins. Molecular recognition elements, found within largely disordered regions, often possess functionally significant residual structure [65] and are key determinants of molecular recognition [66]. Structural capacitance may play a related role in eukaryotic pathogens. Unicellular eukaryotes have considerable variability in their disorder content, which appears to reflect their habitats [18]. The proteomes of parasitic host-changing protozoa, for example, have high levels of disorder, which may represent an adaption to the parasitic lifestyle [67]. Organisms inhabiting environments with high intrinsic frequencies of change, such as microbes, maintain a larger pool of disorder. This is consistent with structural capacitance as constituting a general mechanism for furnishing the capacity to adapt to rapidly changing environments over compressed timescales.

#### Ribosome evolution

We suggest that the D**→**O transition is a necessary consequence of early codon-anticodon usage. The stability of codon:anticodon interactions is determined by the stacking energies of adjacent nucleotides [68, 69]. Wolfenden and colleagues recently postulated that tRNA acceptor stem amino acid recognition is correlated with the size of the amino acid and that anticodon recognition is based on polarity with a strong correlation between polar/non polar and Pu/Py frequency at the 2nd anticodon position [70, 71]. In the prebiotic world the most stable codon:anticodon pairs (most likely a two base pair code initially) would have been selected for the restricted set of amino acids likely to have been present (Ala, Val, Thr, Gly, Asp, and Glu). This would predict an inverse correlation between codon:anticodon stability and amino acid complexity. Consequently, most early proteins would have been disordered or poorly structured. Eventually a critical threshold of amino acids would have been reached, at which point moving to more complex protein structures would be limited by the lack of physicochemically diverse amino acids. Catalysis is required for efficient replication and, in the RNA world this would have arisen from RNA secondary structure. A current hypothesis is that acceptor stem recognition preceded anticodon recognition [70] and that this selection, based essentially on size discrimination led to a prevalence of beta structures with alternating small and large amino acids. However, this argument does not address the question of how these amino acetylated tRNA molecules then assemble on an mRNA template in the absence of anticodons. Irresepective of whichever mechanism originates first - stem recognition or codon recognition - the limited subset of essentially polar amino acids originally present and occupying necessarily the most stable codon:anticodon arrangements supports the notion that the first primordial proteins (maybe as short as 8 amino acids) were relatively poorly structured.

It follows therefore that the D**→**O transition is implicit in the genetic code simply because all the stable codon:anticodon interactions were initially assigned to simple polar amino acids producing little structure. Subsequent mutations would invariably result in a less stable codon:anticodon interaction allowing other amino acids to fill the void. Since other amino acids with few exceptions are physicochemically more complex this would allow for successively increasing amounts of order to appear in protein structure. Finally, the initial set of proteins resulting from this process and comprising the phenotype of primitive organisms would already have a set of structure/function motifs within its proteome. Subsequent O**→**D transitions are likely consequently to be deleterious; D**→**O transitions may still occur, but would require transitioning from stable to unstable codon:anticodon interactions. Since this interaction, is no longer critical with regard to codon usage as the ribosome already exists, the effect would be restricted to protein function, and result in structural capacitance. So the initial evolutionary pressure to select for codon use (namely stable codon:anticodon interactions) is no longer present, and a consequence of the prior pattern of evolutionary change is that all D**→**O mutations which mostly involve stable to unstable codon:anticodon are refractive to subsequent evolutionary modification, except if that pressure is exerted at the level of the ensuing new fold or structure. The codon change involved in D**→**O transitions is neutral as far as selection at the level of codon usage is concerned. Once a primitive ribosome has started to assign codon usage under the exclusive selective pressure of codon:anticodon stability, there is no reversibility. It is remarkable that codon:anticodon stability favors simple amino acids. Structural capacitance is therefore is an invariable consequence of ribosomal evolution.

## Conclusions

The generation of novel microstructures from conformational ‘noise’ through the mechanism of structural capacitance may have contributed to the functional diversification of the protein repertoire through the origin of new ancestral folds, and in so doing contributed to the origin of life and its subsequent elaboration. Although the reported mutations discussed here are associated with diseases representing a number of different pathogenic types including metabolic, vascular, neoplastic and congenital, the subset of O**→**D and D**→**O mutations appears more likely to have a significant causal role and to be ‘drivers’, than O**→**O and D**→**D mutations in which there is no accompanying loss or gain of microstructural change. In summary, the phenomenon of structural capacitance has implications ranging from the ancestral diversification of protein folds to the engineering of synthetic proteins with enhanced evolvability.

## Materials and Methods

The overall workflow is shown in Figure S1.

### Datasets

The target dataset was ‘Human polymorphisms and disease mutations dataset (http://www.uniprot.org/docs/humsavar)[14]. The release of this dataset, according to the UniProt database, is 2 June 2017. This contains 76,608 human mutations including 29,529 human disease mutations, 39,779 polymorphisms and 7,300 unclassified mutations. Disease mutations were annotated using a basic description of the diseases and their OMIM (http://www.ncbi.nlm.nih.gov/omim/) accession number. Disease mutations are labeled based on literature reports. The UniProt database does not systematically annotate mutations as germline or somatic. For each mutation, this dataset provides detailed information including UniProt (http://www.uniprot.org/) accession number of the original protein, mutated position, wild type and mutated amino acid and mutation type (i.e., ‘disease’, ‘polymorphism’ and ‘unclassified’). For the analysis of disease-and non-disease mutations in this study, ‘unclassified’ mutations were removed as the disease annotations for such mutations were ambiguous. Such mutations remain only in Tables S11 and S12 for providing a comprehensive and complete D**→**O candidate list with experimentally verified disordered regions. After removing the sequences containing uncommon amino acids, the resulting dataset contains 12,738 proteins with 68,383 unique single point mutations (28,662 disease-associated mutations, 39,721 polymorphisms).

### Methods

#### Databases/predictors for disordered region prediction

For both wild-type and mutated proteins, the disorder prediction results were defined using 4 predictors, namely: VSL2B [23], IUPred (short and long versions) [72, 73] and DynaMine [74].

##### D2P2 database

D2P2 [75] is an online knowledgebase for protein disordered regions prediction results using nine tools for protein disorder prediction: PONDR VLXT [13], PONDR VSL2B [23], IUPred (short and long versions) [72, 73], Espritz-D [2], Espritz-X [2], Espritz-N [2], PrDOS [3] and PV2 [4]. In addition, in the updated version of D2P2, MoRF regions (predicted by ANCHOR [76, 77]) and post-translational modification sites annotation were used for the investigation of protein binding and function within the disordered regions.

##### DisProt

DisProt (http://www.disprot.org/index.php [28]; Version: 7 v0.3) harbors experimentally verified intrinsically disorder proteins and disordered regions. DisProt provides detailed function classification, function description and experimental evidence for each entry in this database. The advantage of this database is that the disordered regions harbored in DisProt have been experimentally verified. We used the DisProt database to locate mutations that are located in experimentally-validated regions of disorder. We then applied four predictors (VSL2B, IUPred-L, IUPred-S and DynaMine) to predict disorder-order transitions.

##### IUPred

IUPred (http://iupred.enzim.hu/ [72, 73]) maintains two versions of IUPred including IUPred-S and IUPred-L. Here, ‘S’ and ‘L’ refer to the long LDRs and SDRs, respectively. For the ‘S’ option, the model was trained using a dataset corresponding to missing residues in the protein structures. These residues are absent from the protein structures due to missing electron density in the corresponding X-ray crystal structures. These disordered regions are usually short. Conversely for the ‘L’ option, the dataset used to train models corresponds to long disordered regions that are validated by various experimental techniques. In our study, residues with predicted scores equal to or above 0.5 were considered to be located in disordered regions.

##### PONDR-VSL2B

VSL2B [23] is a widely used sequence-based predictor for intrinsically disordered regions, using Support Vector Machine (SVM) [76]. Residues with predicted scores equal to or above 0.5 are considered to be disordered.

##### DynaMine

DynaMine [74], which is trained with a curated nuclear magnetic resonance (NMR) dataset, was used to predict protein disordered regions with only sequence information as the input. Residues with predicted scores ≤0.69 are considered to be located in disordered regions, while those with scores ≥0.8 are predicted to be in the structured regions.

#### Amino acid hydrophobicity indices

Three indices were chosen in our study: Hopp-Woods hydrophilicity index [25], Kyte-Doolittle hydropathy index [26] and Eisenberg hydrophobicity index [24].

#### Predictor for protein transmembrane helices prediction

TMHMM[78] employs hidden Markov model for membrane protein topology prediction. Given the fact the protein transmembrane domains are structurally stable and ordered, TMHMM was used to further validate the predicted disordered regions. Mutations predicted to be in transmembrane regions were discarded.

#### Protein structure BLAST

In order to ensure that wild-type proteins with predicted disordered regions that lack experimentally-determined structures or homologue structures, we performed a BLAST search against the PDB database (http://www.rcsb.org/pdb/software/rest.do) using the protein sequences (e-value cutoff = 0.01). Any proteins with predicted disordered regions and BLAST hits against the PDB database were removed.

#### ELM database mapping

We mapped both disease and non-disease mutations in D**→**O, O**→**D, D**→**D and O**→**O transitions using the ELM (Version: 1.4) (Eukaryotic Linear Motif; http://elm.eu.org/search/ [20]) database. Eukaryotic linear motifs are short linear motifs in eukaryotic proteins, which are predominantly functional modules found in intrinsically disordered regions [21]. Tables S3-S6 show the mapping results of our mutations of both disease and non-disease for the four structural transitions. All ELMs listed in Tables S3-S6 are experimentally verified (i.e., annotated with experimental evidence showing this ELM to be functional.

#### Pfam database mapping

In order to characterise the domain context of mutations, we mapped both disease and non-disease mutations to the Pfam (Release: 30) database [12]. The mapping results for disease and non-disease mutations of D**→**O and O**→**D transitions are shown in Tables S7 and S8.

#### MEROPS database mapping

<u>To extract the</u> proteolytic cleavage sites for the candidates in Table S2, we mapped the protein and mutation information to the MEROPS database [22]. The mapping results for D**→**O mutations in Table S2 are shown in Table S9.

## Author Contributions

CL, LVTC and RZ performed data analysis. BTP, JM, IK, NAB, JS and MB contributed to writing and conceptual advances. GIW assisted in data analysis. AMB, CL and AW designed the study, and drafted the paper.

## Acknowledgements

CL was supported by China Scholarship Council (CSC) and Monash University Joint PhD Student Scholarship (2011630031), and the Bridging Postdoctoral Fellowship of Faculty of Medicine, Nursing and Health Sciences, Monash University (BPF17-0021). LVTC was supported by a Commonwealth Serum Laboratories (CSL)/Undergraduate Research Opportunities Program (UROP) stipend. GIW is an Australian Research Council (ARC) Discovery Outstanding Researcher Award Fellow (DP140100087). NAB is funded by an ARC Future Fellowship (110100223). AMB acknowledges support from the National Health and Medical Research Council (1022688). We thank Dr Morihiro Hayashida and Prof Tatsuya Akustu from Kyoto University for assistance performing the BLAST analysis. We thank Steve Androulakis and the Monash eResearch Centre for assistance with computational tasks, and Mikael Oliveberg, Renwick Dobson, Colin Jackson and Daniel Christ, for helpful comments.

